# Benchmarking foundation models for splice site and exon annotation

**DOI:** 10.64898/2026.02.22.707219

**Authors:** Zitong He, Liliana Florea

**Affiliations:** Department of Computer Science, Johns Hopkins University, Baltimore, MD 21205; Department of Genetic Medicine, Johns Hopkins University, Baltimore, MD 21205

**Keywords:** Benchmarking, Foundation Models, Exon, Splice Site

## Abstract

Recent foundation and deep learning models have brought a generational leap in improving the quality of genome annotation, particularly in identifying genes and their structural elements, including exons and splice sites. However, they are trained on reduced datasets that may not capture biological complexity, such as differences between coding versus non-coding, terminal versus internal, constitutive versus alternatively spliced, and transposable element (TE)-derived exons. We evaluate several foundation models for gene and splice site annotation, including the transformer-based SegmentNT, Enformer and Borzoi, coupled with a segmentation head for per-base resolution, and the CNN-based SpliceAI and AlphaGenome, along with a newly developed fine-tuned model, STEP2h, on different classes of gene elements as described above. We found that the performance of all methods is highest for the class of exons found in their training data class and decreases drastically for classes of exons poorly represented. In particular, performance is highest for protein-coding genes, coding exons, and constitutive exons, and decreases drastically by up to 2-4 fold for non-coding internal exons, terminal exons, and exons that undergo alternative splicing. Similarly, performance is impaired on LINE-1 and *Alu*-derived exons. In contrast, a locally developed CNN model fine-tuned on a specialized TE-exon dataset showed improved performance in this category. Our study highlights the outstanding challenges in gene and exon annotation when leveraging powerful foundation models, and the need for further fine-tuning on judiciously selected classes of data or task-specific models to capture a broader, more diverse spectrum of gene features.

## 1 Introduction

Gene annotation, the process of identifying the structure of genes and their RNA transcripts along the genome, is one of the first and fundamental problems in computational biology. Identifying the genes and their variations in a newly sequenced organism provides a foundation for understanding its biology and functions, which is crucial for scientific breakthroughs in evolution, medicine, and synthetic biology.

Traditional gene annotation methods were developed to determine the exonintron structure of genes in a newly sequenced species. Based on the approach and type of information used, they fall into two categories. Machine learning methods, primarily HMMs and IMMs [17, 13], learned patterns embedded in the sequences of coding, 5’ UTR and 3’ UTR exons, and motifs around the splice sites. In contrast, evidence-based methods leverage gene information in RNA sequencing reads to map them to their genomic locations and then assemble them into gene models [6].

Recently, a spate of powerful deep learning models has been developed to tackle the annotation of genes and other functional elements encoded in the genome, in particular in the context of predicting consequences of splice-altering mutations that lead to the activation of cryptic splice sites, leading to new and potentially pathogenic exons [2]. Convolutional Neural Network (CNN)-based models such as SpliceAI [11] or AlphaGenome [4] use layers to learn hierarchical features from the data, and can identify patterns in DNA or protein sequences leading to the prediction of splice sites and other gene structure elements, including exons, introns, coding sequences, and 5’ and 3’ UTRs. More general DNA foundation models, such as SegmentNT [1] and HyenaDNA [15], capture sequence characteristics across large windows and can tackle multiple annotation tasks simultaneously using different segmentation heads. Lastly, long-range transformer-based architectures such as Enformer [3] and Borzoi [12], although developed to recognize transcription regulatory sequences, including enhancers and promoters, can be retrained to perform fine-grained structural annotation, producing base-level predictions for exons and splice sites when paired with an appropriate segmentation head [1].

These methods have marked a generational leap in genome annotation. However, they are trained, and their performance is measured, on reduced sets of reference annotations that comprise only the very high-confidence features, such as the primary or longest transcript of a gene [11], which do not fully reflect biological complexity. For example, exons undergoing alternative splicing have weak splice signals that modulate the exon’s inclusion in conjunction with the combinatorial action of numerous splicing activator and repressor proteins. Those in non-coding RNA genes and terminal exons do not conform to the codon composition patterns of coding sequences. Lastly, exons generated by exonization of transposable elements, such as the *Alu* and LINE-1 sequences richly represented in the human genome, have historically confounded gene finding methods, leading to excessive false positives at repeat locations across the genome [17].

Benchmarking of these methods is typically reported at the gene level, not distinguishing among classes of gene elements. Moreover, results are often not fully reproducible in independent studies and on new datasets. Hence, we set out to evaluate the methods while differentiating by the feature categories described above, to understand their strengths and limitations. We test a CNN model, SpliceAI, and four foundation models, including AlphaGenome, SegmentNT, SegmentEnformer and SegmentBorzoi, the latter using the backbone of the original Enformer and Borzoi architectures and coupled with a segmentation head to predict splice sites and exons at nucleotide-level resolution. Additionally, we developed a task-specific method, STEP2h (<u>S</u>pliced-in <u>T</u>ransposable <u>E</u>lement <u>P</u>rediction), by combining the SpliceAI dilated convolutional backbone with a segmentation head, and trained it on an extended collection of gene features. We then evaluated all methods across different classes of gene elements for their ability to predict splice sites and exons.

## 2 Results

### 2.1 Description of Models

We briefly describe the technical underpinnings of each model. Additional design details are listed in **Table 1**.

**Table 1.**
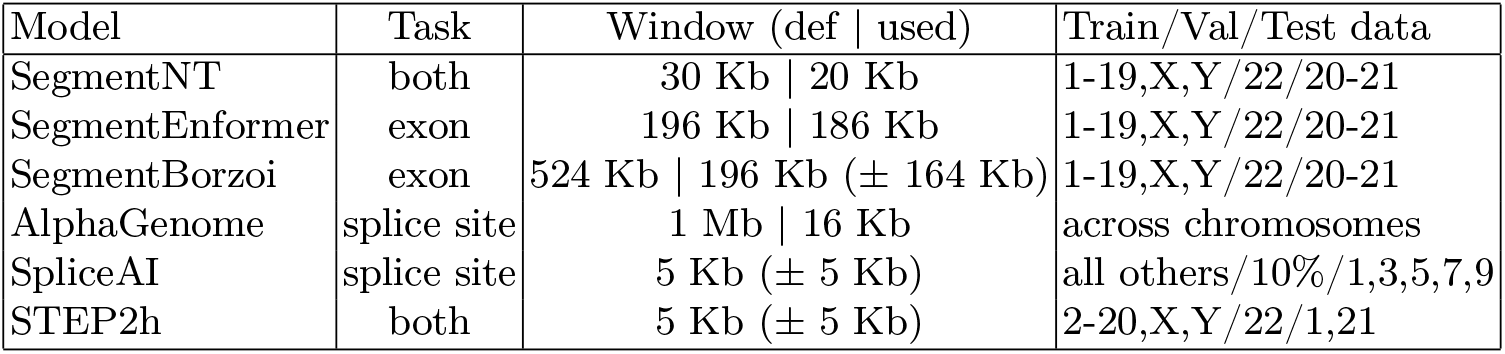
Salient features of models.

**SegmentNT** is a sequence segmentation model built on the 500M-parameter Nucleotide Transformer v2 multi-species encoder, equipped with a 1D U-Net segmentation head that predicts 14 genomic element classes at single-nucleotide resolution, including exons, introns, 5’ and 3’ UTRs, splice acceptors and donors, polyA signals, promoters, enhancers and CTCF sites [1]. In our benchmark, we collapse its 14-way output into: (i) a binary ‘exon’ versus ‘non-exon’ label for the exon segmentation task, and (ii) a three-way splice site view (‘acceptor’, ‘donor’, ‘other’) for splice site detection.

**SegmentEnformer** shares the same 14-class segmentation task and label set as SegmentNT but replaces the backbone with Enformer [3], a convolution–transformer architecture originally developed for long-range regulatory region prediction. The SegmentEnformer variant takes one-hot encoded genome sequence windows of length 196,608 bp (the native Enformer receptive field), and is trained on the same human genome segmentation dataset and chromosome split as SegmentNT.

**SegmentBorzoi** is the analogous segmentation model built on the Borzoi back-bone [12], which was originally trained to predict base-resolution regulatory and RNA-seq profiles from long DNA windows. SegmentBorzoi replaces Bor-zoi’s original task-specific heads with the same 1D U-Net segmentation head as SegmentNT and is trained on the same 14 genomic element classes. It operates on one-hot encoded 524,288 bp windows (the original Borzoi input length) and output 196,608 bp sequence.

**SpliceAI** [11] is a deep dilated CNN designed specifically to predict splice junctions from pre-mRNA sequence. It takes as input long pre-mRNA sequences and, for each nucleotide, outputs the probability that the position is a splice acceptor, splice donor or neither, thereby providing a per-base splice site probability landscape rather than full exon segments.

**STEP2h** is a SpliceAI-like dual-head architecture developed in house, which bridges exon segmentation and splice site detection within a single compact model. It uses the same basic residual dilated-convolution backbone as SpliceAI but doubles the number of channels per layer, increasing capacity while keeping the receptive field comparable. On top of the shared backbone, we add two task-specific heads. The first is an exon segmentation head that outputs per-base probabilities for ‘exon’ versus ‘non-exon’ labels, allowing us to use the model directly in the exon segmentation task. The second is a splice site head that outputs, for every position, probabilities for ‘acceptor’, ‘donor’, and ‘other’ site, mirroring the label space of SpliceAI.

For training, we first fit the model on GENCODE gene sequences to learn the general exon–intron structure and canonical splice sites, and then fine-tune it on TE–derived exons to sharpen its performance on this exon class. Because both exon boundaries and splice sites are highly imbalanced relative to background bases, we train both heads with a focal loss, which downweights easy negatives and focuses optimization on the rare but informative positive positions. This design allows us to evaluate a single, task-specific model that is jointly optimized for exon segmentation and splice site detection, and to test whether explicit TE-aware fine-tuning helps close the gap between TE-derived and non-TE exons.

**AlphaGenome** [4] is a large multimodal DNA foundation model that takes up to 1 Mb of human or mouse genomic sequence as input and predicts thousands of molecular readouts at single-base resolution, including gene boundaries, RNA expression levels, 3D chromatin contacts and, importantly for this work, splice junction locations and usage across many tissues and cell types. Its architecture combines convolutional layers to detect local sequence motifs with transformers to propagate information across the full 1 Mb window, and is trained on a large collection of public functional genomics datasets from ENCODE, GTEx, the 4D Nucleome project, FANTOM5 and related resources.

We align each model with the task that best matches its native output and training regime. SegmentNT, SegmentEnformer and SegmentBorzoi are evaluated on exon segmentation, where their multi-class segmentation heads provide explicit exon labels over long genomic contexts. SpliceAI and AlphaGenome are evaluated on splice site detection, where they naturally provide per-base acceptor and donor site probabilities. Our SpliceAI-like dual-head model is evaluated on both tasks, since it is explicitly designed and trained to produce exon segments and splice site probabilities jointly, providing a compact, TE-aware alternative to the larger foundation models.

### 2.2 Benchmark Results

We compare the performance of all methods across classes of genes, gene exon collections (‘bag of exons’), and exons. First, we evaluate the performance on gene sets, starting with the *reduced* gene and exon set used for training SpliceAI and the foundation models, followed by the extended *‘bag of exons’* annotation of the same gene set, and by a *comprehensive* gene and exon collection curated from GENCODE. Additionally, we compared the methods’ performance on the *coding* versus *non-coding* genes. Second, we distinguish programs’ behavior across different classes of exons by comparing *constitutive* versus *alternatively spliced* exons, and *terminal* versus *internal coding* and *internal non-coding* exons. Lastly, we evaluate the detection of *TE-derived* exons.

### Performance decreases with ‘unseen’ exons and genes

To assess the ability of the programs to detect ‘novel’ genes and transcripts, we apply them to: i) the reduced dataset consisting of one representative transcript for each of 18,475 genes used for training SpliceAI and, with minor variations, all of the published models; ii) the extended ‘bag of exons’ annotation of the same gene set; and iii) a comprehensive annotation dataset consisting of curated ‘bags of exons’ for 35,083 protein coding and lncRNA GENCODE genes (**Fig. 1A**). For each of the three annotations, models performed similarly, with small variations. STEP2h, optimized for exon-segment detection, traded recall for precision at the bitwise level, and SpliceAI consistently outperformed all other models in splice site detection accuracy. Thees trends persisted for the rest of the analyses. When comparing across datasets, all models’ performance was highest on the reduced dataset (avg. F1_bw_ = 0.756, F1_seg_ = 0.810), and decreased with the extended exon set (avg. F1_bw_ = 0.660, F1_seg_ = 0.724), and even further on the comprehensive gene and exon annotation (avg. F1_bw_ = 0.624, F1_seg_ = 0.664). Trends were similar for splice site detection, with average F1 scores of 0.90, 0.83, and 0.75, respectively. The drop in performance was due primarily to lower recall as more exons (20% at segment level and 13% bitwise) and then genes (additional 6% and 5% segment-level and bitwise, respectively) were added. STEP2h, trained on an extensive ‘bag of exons’, exhibited more minor drops (total of 7% drop bitwise, and 14% at segment-level).

**Fig. 1.**
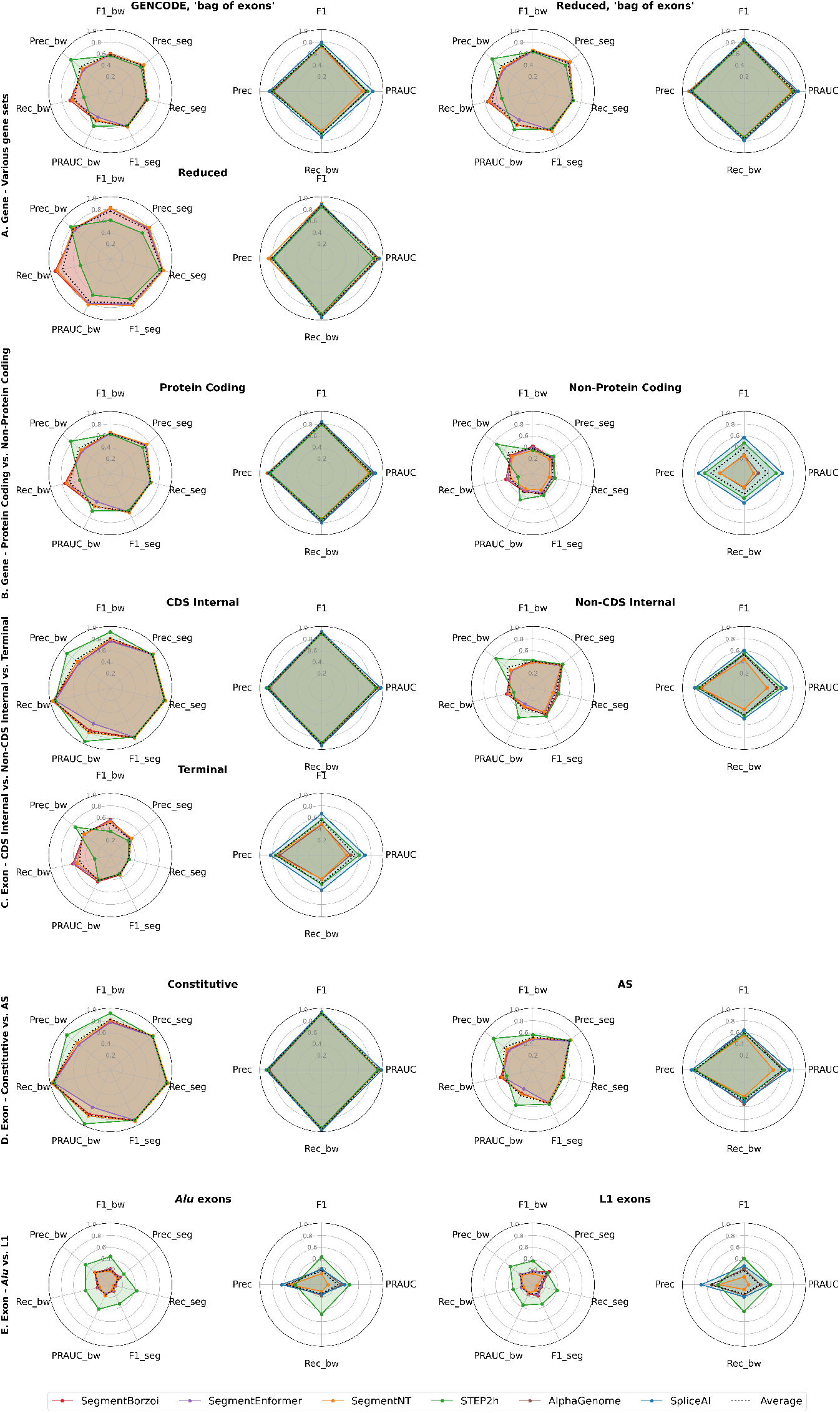
Method performance on classes of genes and exons.

### Performance is lower on non-coding versus protein-coding genes

Protein coding genes contain features that have traditionally been well captured by machine learning methods, such as the codon-based composition and codon bias of ORFs, and start and stop codons, unlike non-coding genes. We assessed the ability of the methods to detect the two classes of genes using the ‘bag of exons’ annotations for 19,045 protein coding and 16,038 non-coding GENCODE genes (**Fig. 1B**). Within each category methods performed similarly, following the trends described above. SpliceAI outperformed all other methods in predicting splice sites, followed by STEP2h. When comparing between classes, methods performed significantly lower on the non-coding class, with an average F1_seg_ of 0.433 versus 0.719 (60% decrease) at the exon level, and 0.378 (0.426) versus 0.817 (0.813) at acceptor (donor) sites, a two-fold decrease. Lower recall rates (by 1.7-2.7 fold) and, to a smaller extent, lower precision contributed to the degradation in performance.

### Performance is lower on terminal and internal non-coding exons compared to coding exons

Sequences of terminal, internal coding and internal non-coding exons undergo different selective pressures and differ in their length distributions, which have traditionally required different machine learning models [17]. Next, we evaluated the methods on the 132,861 terminal, 170,560 internal-coding, and 84,268 internal-noncoding exons extracted from the GENCODE ‘bag of exons’ annotation (**Fig. 1C**). Methods’ performance is highest on coding (average F1_seg_ 0.895), followed by internal non-coding (0.511), and terminal (0.381) exons – a 2.3 fold drop. For splice sites, accuracy as measured by F1 is highest for splice site of coding exons (0.893 acceptor | 0.899 donor), followed by terminal (0.577 | 0.566) and internal non-coding (0.497 | 0.552) exons, pointing to the interplay between the strength of splicing regulatory signals and exonic sequence composition in shaping the methods’ power to recognize features. Both lower recall and lower precision contributed to the exon-level decrease in performance, whereas precision was relatively stable while recall dropped sharply (2.1-2.4 fold, from over 90% to 37-43%).

### Performance is lower on alternatively spliced compared to constitutive exons

Alternatively spliced exons generally harbor weaker splice signals [9], to allow modulation of splicing by regulatory factors that work in a combinatorial set of interactions to determine the splice site selection. We constructed a comprehensive set of 60,495 alternatively spliced (‘skipped’) exons from across CHESS3, MANE, and GENCODE v.46 gene annotations (see Methods), and 116,384 constitutive exons, included in MANE Select transcripts and without evidence of alternative splicing. Methods showed uniformly high accuracy in detecting constitutive exons and their splice sites, with average F1 scores of 0.914, 0.907, and 0.901, respectively. These values decreased by 25-30% for alternatively spliced exons, to 0.626, 0.663, and 0.679; the drop in performance was even more accentuated for SpliceAI, by 32.3%-35.6%. Decreases in recall, by 37.7-42.3%, were the primary driver of performance loss, whereas precision decreased by less than 11% (**Fig. 1D**).

### Performance degrades on TE-derived exons

TE-derived exons have traditionally been challenging to detect with inferential methods. Few examples are present in reference gene annotations for methods to learn patterns from, and when included in the training data, they lead to large numbers of false positives at repeat sites across the genome. Consequently, most annotation pipelines are run on the repeat-masked genome. Further, TE-exons are more likely to be non-coding, alternatively spliced, expressed at low levels, or expressed in a tissue or condition-specific manner [2]. We evaluated methods for identifying 25,180 *Alu* and 7,636 LINE-1 exons obtained from an analysis of GTEx data (see Methods). Conventional methods’ performance was significantly lower than on previously described classes of exons: segment level accuracy only reached 0.15-0.20, marked by drops in both recall (0.11-0.14) and precision (0.26-0.36), whereas splice site detection retained higher precision (0.56-0.60), at low recall (0.13-0.17). Our task-specific model STEP2h, fine-tuned for *Alu* and LINE-1 exons, respectively, significantly improved segment-level performance by increasing recall 3.5-4.5 fold as precision remained stable. At the splice site-level, the same increase in recall was met with a roughly 20% decrease in precision, leading to a 2-fold improvement in the F1 score overall (**Fig. 1E**).

## 3 Methods

### 3.1 Materials and Data

For the *gene-level* evaluation, we used GENCODE v.46 gene annotations. For the coding versus non-coding gene set evaluation, we extracted genes in each category based on the *gene_type* label: ‘protein_coding’ versus ‘lncRNA’. Then, for each gene, we constructed a comprehensive *‘bag of exons’* by aggregating all unique exon intervals across its annotated transcripts, retaining only multi-exon genes, and removing redundant or ambiguous exon boundaries. This produced a unified exon set per gene that reflects the complete exon structure observed across its transcript isoforms. In contrast, the reduced training data used by SpliceAI comprises only the longest transcript of a gene and all protein-coding genes. *Exon* categories are described below and illustrated in **Fig. 2**.

**Fig. 2.**
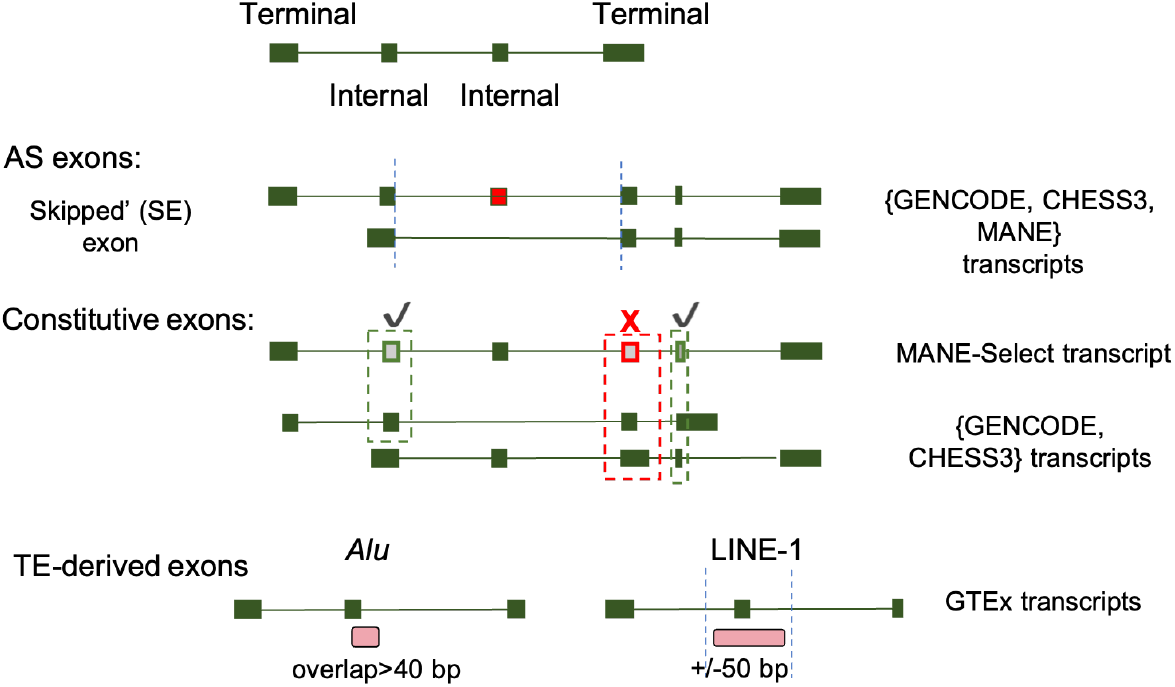
Classes of exons surveyed.

**Fig. 3.**
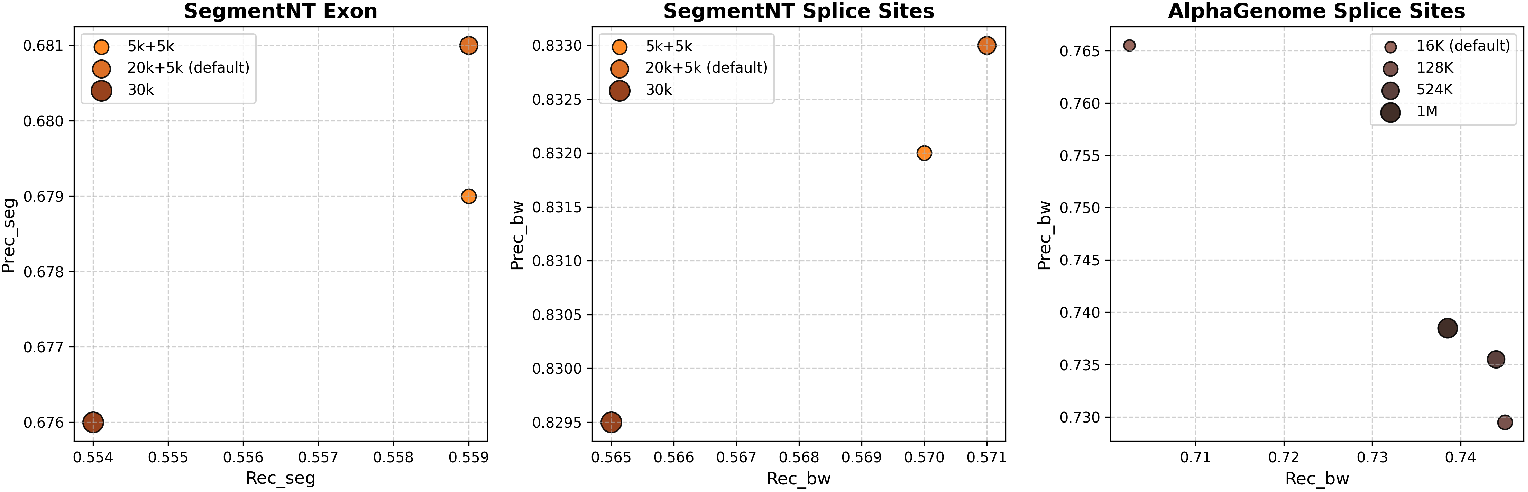
Model input range calibration.

#### Terminal, internal coding, and internal non-coding exons

‘Terminal’ exons refer to the 5’- and 3’-most exons in a transcript. For internal exons, we use the reading frame labels in the genes’ ‘CDS’ records to distinguish coding from non-coding exons.

#### Alternatively spliced and constitutive exons

A ‘constitutive’ exon is traditionally defined as one that occurs in all transcripts of a gene, whereas an ‘alternatively spliced’ exon occurs in only a subset of transcripts. Accurately identifying members of the two classes, however, has become more challenging, as reference databases remain incomplete and newly discovered transcripts may prove some constitutive exons to be alternatively spliced. Further, adding more transcript variants, often computationally inferred and potentially inaccurate or artifactual, to the GENCODE annotation may cause a true constitutive exon to appear as alternatively spliced. We aimed to select a high-confidence collection of constitutive and alternatively spliced exons, using a conservative method. First, we selected all exons designated as ‘skipped’ (‘SKIP_ON’ label) when combining the MANE v.1.4, CHESS3 v.3.1.3, and GENCODE v.46 annotations, as classified with ASprofile [7], as *alternatively spliced*. For *constitutive* exons, we select all internal exons from the MANE-Select database that do not have any overlap with other (internal) exons. MANE-Select represents a high-confidence set of transcripts, on a per gene basis, that are identically annotated between the Ensembl and RefSeq datasets, and have been manually evaluated for accuracy against evidence.

#### Transposable element (TE)-derived exons

We denote a TE-exon as one that has resulted from exonization, or recruitment (fully or partially) of a TE into a gene transcript as a spliced exon. Exons derived from the 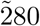 bp *Alu* retrotransposon and from the 2-7Kb *LINE-1* element have contributed the most to exonization. As TE-exons are poorly represented in reference databases including GENCODE [8], we identified *Alu* and LINE-1 exons from analyses of GTEx data in 28 tissues (277 RNA-seq samples) as described in [10]. Briefly, reads were mapped to the human genome GRCh38 using STAR [5] and assembled into transcripts with CLASS2 [16]. Then, internal exons 40-1000 bp long and overlapping *Alu* and LINE-1 genomic annotations by >25 bp were selected as internal TE-exons.

### 3.2 Method Benchmarking

#### Software

We obtained SegmentNT, SegmentEnformer, and SegmentBorzoi from their Hugging Face repositories and ran them locally. AlphaGenome was obtained from its GitHub repository and queried remotely via its CLI interface. SpliceAI was obtained from its GitHub repository and run locally.

#### Calibration

By default, the tested models have vastly different input genomic ranges, motivated in part by their multi-class prediction function. To ensure compatibility among the models, and because splicing interactions are more local, we tested the longer range models on shorter intervals. We tested AlphaGenome’s splice site prediction function on 16 Kb, 124 Kb, 524 Kb and 1Mb sequences, and SegmentNT’s splice site and exon prediction functions on 5 Kb (±5 Kb of context) for the splice, 20 Kb (±5 Kb of context) and 30 Kb. The models’ performance was robust with the input sequence length for the two tasks 3, and we chose the marginally better options, namely 16 Kb input for AlphaGenome and 20 Kb for SegmentNT **(Table 1)**.

#### Model running

For efficiency, we run the tools on the entire chr1 and chr21 in windows as specified by the tool’s protocol. Once the predictions are calculated, we extract and calculate the performance measures in the areas of interest. For evaluation by class of exons, annotations at exons other than the target are “masked” out from the region. Additionally, because exon sets may extend over only a portion of the gene, we scale the ‘background’ to the query exon set, assuming a uniform distribution of the exons’ lengths within the gene span. Specifically, *FP*_*norm*_ = *αFP*, where *α* is the proportion of bases in the target exons to the total size of exons in the reference annotation; all other measures are adjusted accordingly.

#### Test set and measures

Given the tools original training and validation sets, benchmarking all models on held-out chromosomes 1 and 21 provides a fair and diverse genomic test set for our own model - trained on all other chromosomes, excluding chr1 and chr21; and for SegmentNT/SegmentEnformer/SegmentBorzoi - whose original training protocol explicitly holds out chr21 but not chr1; while still allowing a consistent comparison to SpliceAI - which was originally trained on genes including chr21 but tested on chr1; and to AlphaGenome - which was pre-trained genome-wide and may have seen both chromosomes during training. We use TP, FP, TN, FN, 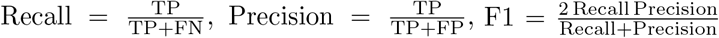, and PRAUC as measures, at the bitwise, splice site, and segment levels. A cutoff of 50% intersection over union (IoU), defined as the overlap between two intervals divided by their combined span, is used to determine a segment-exon match.

## 4 Discussion and Conclusions

We evaluated four foundation and one CNN sequence models recently reported in the literature for their ability to detect splice sites and exons, which are essential to the gene annotation process. We found that methods consistently do not meet the published values, often generated on a reduced dataset, when presented with a full spectrum of features. Specifically, methods perform best and within expectations on protein coding genes, especially the subset of genes included in the reduced training set for SpliceAI and other models, and when only the representative transcript per gene is included. In contrast, accuracy is reduced by half on non-coding genes. Among classes of exons, constitutive exons and coding exons and their splice sites are the most reliably predicted, with ∼0.90 F1 accuracy. Performance is significantly reduced, by up to two times to ∼0.40-0.50 F1 scores, for non-coding internal exons and terminal exons, and to a lesser but significant degree for alternatively spliced exons, to ∼0.60 F1 scores. The prediction is further degraded on transposable element-derived exons, such as *Alu* and LINE-1, which exhibit a greater than 4-fold drop, to the low 0.15-0.3 F1 accuracy measures.

The importance of these discrepancies is significant, particularly in the context of current biomedical applications that seek to identify changes in gene structure associated with diseases. Many elements in the classes highlighted above may have biomedical relevance. For example, transposable element-derived exons can be linked to genetic diseases, and condition specific or alternatively spliced exons, including activated cryptic exons, can occur in cancer and other diseases, such as frontotemporal dementia [14].

The limited training and evaluation datasets used by these methods are significant contributors to performance issues. In our analyses, we observed that considerably lower recall rates were the primary factor behind performance loss, while precision declined more gradually. This limitation could potentially be addressed by incorporating more examples into the training dataset or by recalibrating the programs. Additionally, biological complexity, such as the repetitive nature and prevalence of transposable elements in the genome along with high sequence similarity among occurrences, can also limit performance. For example, both recall and precision were impacted for the TE-exons class. To address these challenges, targeted approaches, starting with a general model and then fine-tuning it for specific exon classes, such as with STEP2h, or independently developed task-specific systems [10], could yield better results.

## Acknowledgments

This work was supported in part by grant R35GM156374 from the National Institutes of Health, and award DBI-2504115 from the National Science Foundation, to L.F..

